# Variations in H_2_ thresholds and growth yields reveal bioenergetic diversity among hydrogenotrophic methanogens

**DOI:** 10.1101/2025.08.26.672303

**Authors:** Timothé Philippon, Jo Philips

## Abstract

Hydrogenotrophic methanogens are of high environmental and biotechnological importance, converting CO_2_ with H_2_ into CH_4_. Despite their common metabolism, variations in the energy metabolism among these methanogens exist, likely affecting their H_2_ thresholds and growth yields. However, a systematic comparison of these traits for a wide range of hydrogenotrophic methanogens has been lacking. Here, we measured the H_2_ thresholds and growth yields of nine different hydrogenotrophic methanogens. The H_2_ threshold, i.e. the H_2_ partial pressure at which H_2_ consumption halts, ranged over two orders of magnitude from 1.0 ± 0.5 Pa for *Methanobrevibacter arboriphilus* to 120 ± 10 Pa for *Methanosarcina mazei*. Growth yields in our experimental conditions ranged from 0.51 ± 0.28 g_DCW_×(mol CH_4_)^−1^ for *Methanococcus maripaludis* to 5.28 ± 1.25 g_DCW_×(mol CH_4_)^−1^ for *Methanosarcina mazei*. The ATP gains, estimated from both H_2_ thresholds and growth yields, correlated reasonably well, confirming that these variations are due to differences in energy conservation strategies. Our results strongly differentiated the two previously proposed groups of hydrogenotrophic methanogens: methanogens with cytochromes had a high H_2_ threshold (≥ 21 Pa) and high growth yield (> 4.0 g_DCW_×(mol CH_4_)^−1^), whereas methanogens without cytochromes had lower H_2_ threshold (≤ 7 Pa) and low growth yield (< 1.7 g_DCW_×(mol CH_4_)^−1^). Moreover, our H_2_ thresholds indicated that additional variations in energy metabolism exist within both groups. Overall, this study found strong variations between hydrogenotrophic methanogens, which are important to understand their environmental prevalence and biotechnological applicability.

**Importance:** Hydrogenotrophic methanogens play key roles in natural ecosystems and biotechnological processes. Even though all hydrogenotrophic methanogens convert CO_2_ with H_2_ into CH_4_, they can differ in their H_2_ threshold and growth yield, due to variations in their energy metabolism. Here, we found that H_2_ thresholds of hydrogenotrophic methanogens range over two orders of magnitude, while a ten-fold difference was observed in their growth yields. These strong variations in H_2_ thresholds and growth yields demonstrate that hydrogenotrophic methanogens are confronted with a trade-off between the capacity to grow at low H_2_ levels (low H_2_ threshold) and efficient growth (high growth yield). Curiously, the observed H_2_ thresholds also correlate with the electroactivity of different hydrogenotrophic methanogens. Overall, by reporting H_2_ thresholds and growth yields for a wide range of hydrogenotrophic methanogens, this work provides new insights into the bioenergetic diversity of these microbes, as well as their environmental prevalence and biotechnological applicability.

## Introduction

Hydrogenotrophic methanogenesis is a metabolic pathway used by specific Archaea to produce methane (CH_4_) from carbon dioxide (CO_2_) and dihydrogen (H_2_) for the purpose of energy conservation:

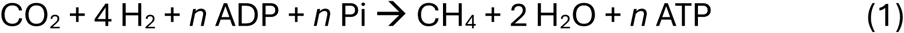

This metabolism is widely spread in the environment and plays an important role in the global carbon cycle, as methanogens convert yearly about one gigaton of carbon into methane [1]. This production of methane is a major issue in the context of climate change, as methane is a potent greenhouse gas. Methanogenesis can also negatively impact steel infrastructure, as certain hydrogenotrophic methanogens induce strong corrosion [2]. Nevertheless, hydrogenotrophic methanogenesis is of huge interest for the development of energy conversion (e.g. power-to-gas) biotechnologies, as methane is a valuable fuel. Bio-methanation is already an advanced technology, converting CO_2_-containing gases into methane with the addition of H_2_ gas (Reaction 1) [3]. Another promising biotechnology is electro-methanogenesis, in which an electrode is used to generate H_2_ *in situ* of a bioreactor [4, 5]. Because of the technological and environmental importance of hydrogenotrophic methanogens, variations between different methanogens need to be well understood.

All hydrogenotrophic methanogens use a common pathway for their energy metabolism, consisting of a succession of steps reducing CO_2_ to CH_4_, using electrons obtained from the oxidation of H_2_ and generating a sodium (Na^+^) gradient (Figure 1) [6]. Besides this common pathway, hydrogenotrophic methanogens vary in the involvement of cytochromes [1, 6]. Hydrogenotrophic methanogens without cytochromes (Figure 1A) generate ATP only through the Na^+^ gradient [7]. In contrast, *M. barkeri* creates an additional H^+^ gradient through two protein complexes (*Vht* and *hrdDE*), each containing cytochromes (Figure 1B) [6]. Methanogens with cytochromes thus have a higher ATP gain (amount of ATP generated (*n*) in Reaction 1) than methanogens without cytochromes (Table 1, Figure 1). Moreover, further variations in the energy conservation components of both groups of methanogens have been reported [6, 8, 9], suggesting that additional differences in ATP gains may exist.

**Figure 1:**
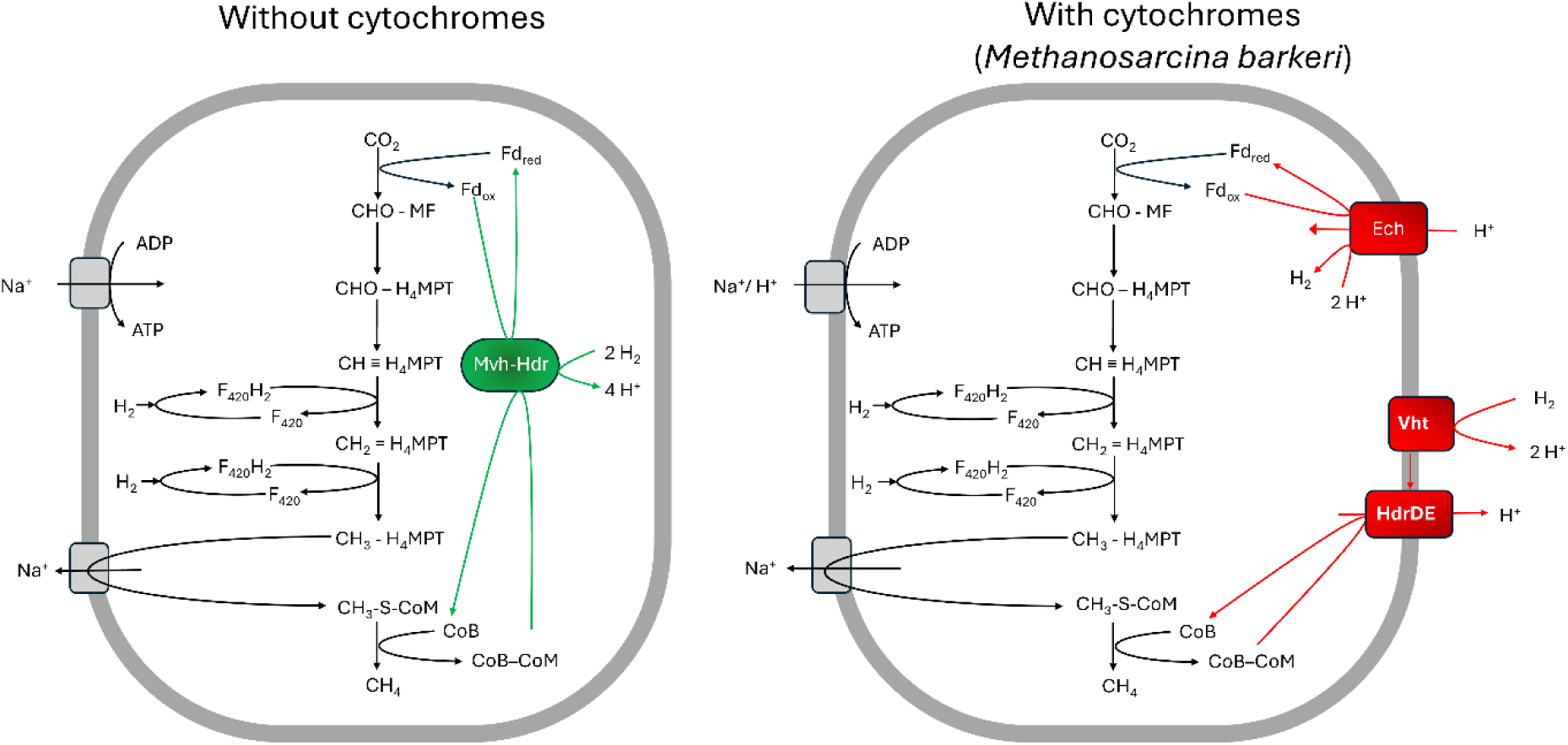
Energy metabolism for hydrogenotrophic methanogenesis without cytochromes (left) and with cytochromes (for *M. barkeri*) (right). Black colored components and reactions are common for all hydrogenotrophic methanogens, including the generation of a sodium (Na^+^) gradient through the transfer of the methyl group from tetrahydrometanopterin (H_4_PMT) to Coenzyme M (CoM). Hydrogenotrophic methanogens without cytochromes generate ATP only through this Na^+^ gradient. They use flavin-based electron bifurcation to couple the endergonic reduction of Ferredoxin (Fd) with H_2_ as electron donor to the exergonic reduction of CoM-CoB, through the action of the *Mvh-Hdr* ([NiFe]-hydrogenase - heterodisulfide reductase) complex. In contrast, *M. barkeri* (with cytochromes) generates ATP through both a Na^+^ and H^+^ gradient. In addition to the Na^+^ gradient, *M. barkeri* creates a H^+^ gradient through two protein complexes *Vht* (methanophenazine-reducing hydrogenase) and *hrdDE* (methanophazine-dependent heterodisulfide reductase), each containing cytochromes and together forming an electron transport chain transporting electrons from H_2_ to CoM-CoB and pumping H^+^ out of the cell. In addition, *M. barkeri* reduces Fd with the electron donor H_2_ through consumption of the H^+^ gradient using an *Ech* (energy converting hydrogenase) complex. Other abbreviations: methanofuran (MF), Coenzyme B (CoB)[6, 8].

**Table 1:** Comparison of metabolic characteristics of hydrogenotrophic methanogens depending on the presence or absence of cytochromes according to previous literature, as well as this study.

|  | Methanogens without<br>cytochromes | Methanogens with<br>cytochromes |
| --- | --- | --- |
|  | Including<br>*Methanococcus*<br>*Methanobrevibacter*<br>*Methanobacterium*<br>*Methanolacinia* | Including<br>*Methanosarcina* |
| H<sub>2</sub> threshold (Pa) |  |  |
| Previous literature (Table S1) | ≤20 | ≥13 |
| This study (Table 2) | ≤7.0 | ≥21 |
| Growth yield (g<sub>DCW</sub> · mol<sup>-1</sup> CH<sub>4</sub>) |  |  |
| Previous literature (Table S2) | 0.8 - 3.5 | 5.4 - 8.4 |
| This study (Table 2) | 0.5 - 1.7 | 4.1 - 5.3 |
| ATP gain (mol ATP · mol<sup>-1</sup> CH<sub>4</sub>) |  |  |
| [7] | <1 | >1 |
| This study, based on H<sub>2</sub> threshold (Table 2) | 0.32-0.56 | 0.75-0.95 |
| This study, based on growth yield (Table 2) | 0.10-0.34 | 0.82-1.06 |

Different ATP gains in turn affect various metabolic characteristics of methanogens, including their H_2_ threshold and growth yield (Table 1). The H_2_ threshold quantifies the H_2_ concentration at which H_2_ oxidation and therefore methanogenesis halts due to thermodynamic constraints. This H_2_ threshold is generally lower for methanogens without cytochromes than for methanogens with cytochromes (Table 1 and S1). Accordingly, the growth yield, i.e. the amount of biomass created per amount of CH_4_ produced, is higher for methanogens with cytochromes than for methanogens without cytochromes (Table 1, Table S2). Additional variations in these metabolic characteristics likely exist, but a coherent comparison of an extensive number of methanogens has not yet been performed.

Most studies measuring growth yields for hydrogenotrophic methanogens included just one to two methanogenic species, while each using different media and experimental setups (Table S2). However, growth yields strongly depend on the medium composition [10, 11], the gas supply [12], the dissolved H_2_ concentration [13] and the growth rate [14] (Table S2). Growth yields determined in different studies are thus not directly comparable. Similarly, most studies measuring H_2_ thresholds included only few methanogens, each using different conditions (Table S1). H_2_ thresholds depend on the experimental conditions (e.g. temperature, total pressure, CO_2_ and CH_4_ levels and even trace element concentrations [15–17]), which likely explains why reported H_2_ thresholds for e.g. *Methanosarcina barkeri* differ widely (from 13 to 150 Pa, Table S1). H_2_ thresholds determined in different studies are thus neither directly comparable. Recently, strongly different H_2_ thresholds were unexpectedly found for acetogens [18]. It seems thus worthwhile to revisit the H_2_ thresholds and growth yields of different methanogens.

Insights into the H_2_ thresholds of different methanogens are also crucial to assess their potential for electromethanogenesis and role in microbial-induced corrosion. In both cases, the cathode or steel surface acts as extracellular electron donor, from which methanogens withdraw electrons, using H_2_ as main intermediate [5]. Not all hydrogenotrophic methanogens are as effective in this extracellular electron uptake [19]. Besides the involvement of extracellular hydrogenases [20], the H_2_ partial pressure is proposed to play a role in electroactivity [15]. Microbes maintaining low H_2_ partial pressures likely favor H_2_ evolution from a cathode or steel surface. As the H_2_ threshold reflects the lowest H_2_ partial pressure a strain can sustain, this parameter is thus likely crucial to understand the electroactivity of methanogens.

This study aimed to compare H_2_ thresholds and growth yields of nine strains of mesophilic hydrogenotrophic methanogens in similar conditions (Table 2). The selected methanogens include species with and without cytochromes, originating from different environments and spanning various methanogenic genera. In addition, two methanogenic strains known for their strong corrosive behavior were included. For all these strains, the H_2_ threshold and the growth yield were experimentally measured. Moreover, ATP gains were estimated from either the H_2_ threshold or the growth yield. All these variables are relevant to assess variations among methanogens.

**Table 2:**
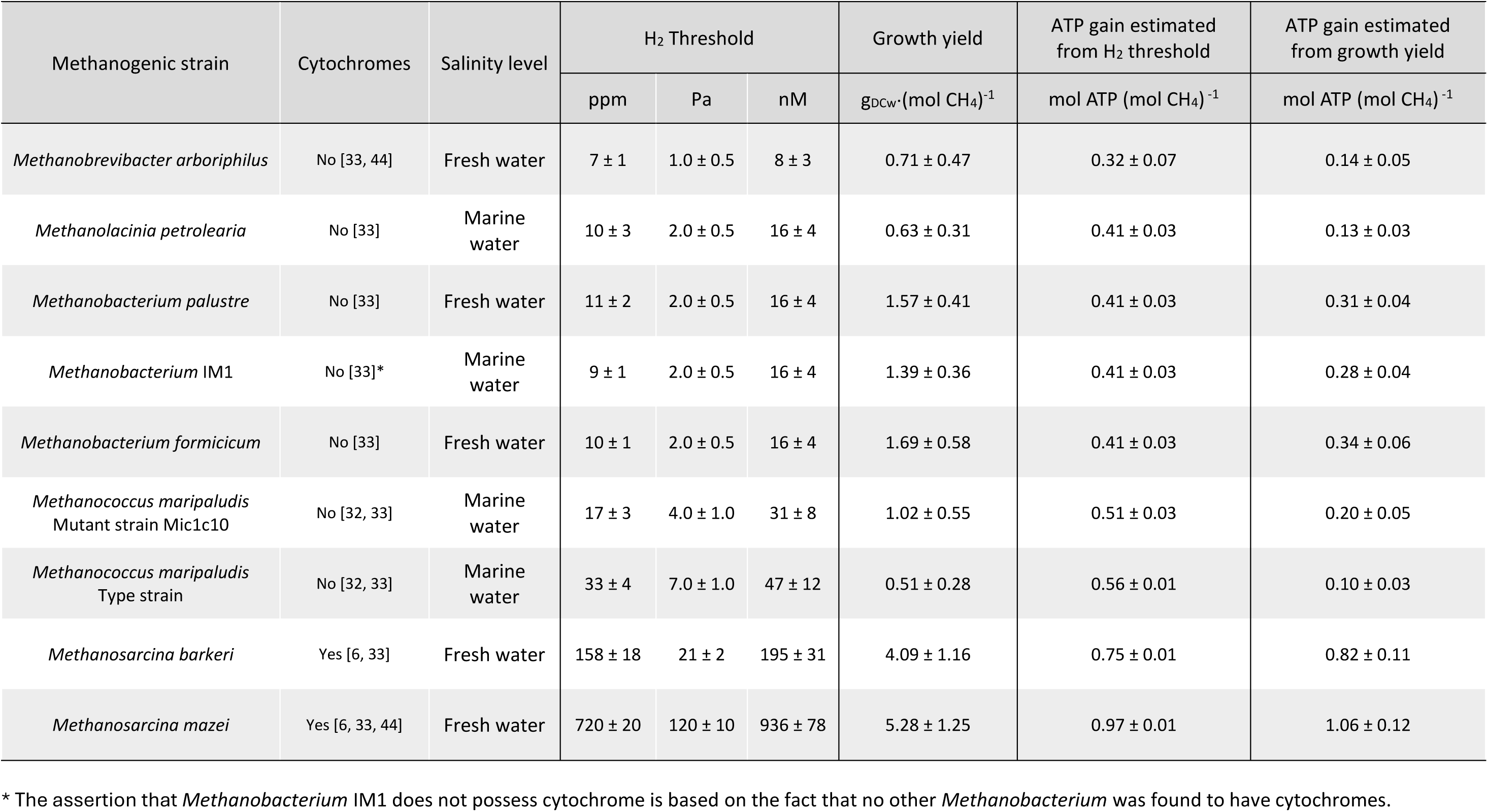
Overview of the methanogenic strains examined in this study, as well as their experimentally determined H_2_ thresholds andgrowth yields. The table also includes the ATP gains estimated either from the experimental H_2_ thresholds or the experimental growth yields. The H_2_ thresholds measured are expressed as partial pressures in the headspace (both in ppm and Pa) and as concentration dissolved in the liquid phase (nM).

| Methanogenic strain | Cytochromes | Salinity level | H<sub>2</sub> Threshold |  |  | Growth yield<br>g<sub>DCW</sub>·(mol CH<sub>4</sub>)<sup>-1</sup> | ATP gain estimated<br>from H<sub>2</sub> threshold<br>mol ATP (mol CH<sub>4</sub>)<sup>-1</sup> | ATP gain estimated<br>from growth yield<br>mol ATP (mol CH<sub>4</sub>)<sup>-1</sup> |
| --- | --- | --- | --- | --- | --- | --- | --- | --- |
|  |  |  | ppm | Pa | nM |  |  |  |
| *Methanobrevibacter arboriphilus* | No [33, 44] | Fresh water | 7 ± 1 | 1.0 ± 0.5 | 8 ± 3 | 0.71 ± 0.47 | 0.32 ± 0.07 | 0.14 ± 0.05 |
| *Methanolacinia petrolearia* | No [33] | Marine water | 10 ± 3 | 2.0 ± 0.5 | 16 ± 4 | 0.63 ± 0.31 | 0.41 ± 0.03 | 0.13 ± 0.03 |
| *Methanobacterium palustre* | No [33] | Fresh water | 11 ± 2 | 2.0 ± 0.5 | 16 ± 4 | 1.57 ± 0.41 | 0.41 ± 0.03 | 0.31 ± 0.04 |
| *Methanobacterium* IM1 | No [33]* | Marine water | 9 ± 1 | 2.0 ± 0.5 | 16 ± 4 | 1.39 ± 0.36 | 0.41 ± 0.03 | 0.28 ± 0.04 |
| *Methanobacterium formicicum* | No [33] | Fresh water | 10 ± 1 | 2.0 ± 0.5 | 16 ± 4 | 1.69 ± 0.58 | 0.41 ± 0.03 | 0.34 ± 0.06 |
| *Methanococcus maripaludis*<br>Mutant strain Mic1c10 | No [32, 33] | Marine water | 17 ± 3 | 4.0 ± 1.0 | 31 ± 8 | 1.02 ± 0.55 | 0.51 ± 0.03 | 0.20 ± 0.05 |
| *Methanococcus maripaludis*<br>Type strain | No [32, 33] | Marine water | 33 ± 4 | 7.0 ± 1.0 | 47 ± 12 | 0.51 ± 0.28 | 0.56 ± 0.01 | 0.10 ± 0.03 |
| *Methanosarcina barkeri* | Yes [6, 33] | Fresh water | 158 ± 18 | 21 ± 2 | 195 ± 31 | 4.09 ± 1.16 | 0.75 ± 0.01 | 0.82 ± 0.11 |
| *Methanosarcina mazei* | Yes [6, 33, 44] | Fresh water | 720 ± 20 | 120 ± 10 | 936 ± 78 | 5.28 ± 1.25 | 0.97 ± 0.01 | 1.06 ± 0.12 |

\* The assertion that *Methanobacterium* IM1 does not possess cytochrome is based on the fact that no other *Methanobacterium* was found to have cytochromes.

## Material and methods

### Methanogenic strains and culture conditions

This study included nine different methanogens (Table 2). Seven methanogenic strains were ordered from the Leibniz Institute DSMZ, including *Methanococcus maripaludis* (DSMZ 2067 designated as type strain in this study), *Methanolacinia petrolearia* (DSMZ 11571), *Methanobacterium palustre* (DSMZ 3108), *Methanobacterium formicicum* (DSMZ 1535), *Methanosarcina mazei* (DSMZ 2053), *Methanosarcina barkeri* (DSMZ 800) and *Methanobrevibacter arboriphilus* (DSMZ 744). Once received, the strains were transferred to their recommended DSMZ medium (exact composition but without Na_2_S, sludge fluid or fatty acid mixture) and stored frozen at – 80 °C in a 2 mL tube with 25% glycerol.

In addition, a culture containing *Methanobacterium* strain IM1 [21, 22] was obtained from Florin Musat (Aarhus University). The culture was inoculated in DSMZ medium 141 (without Na_2_S) and was stored frozen at – 80 °C. Microscope imaging (Leica ICC50 W, Germany) showed that an additional coccus shaped microbe was present in this culture. Nevertheless, when cultivated in a medium with only H_2_ as electron donor (no acetate), the IM1 strain (bacille shape [22]) became dominant. Acetate was thus further omitted from the medium. Therefore, we assume that the measured H_2_ threshold was for *Methanobacterium* strain IM1.

*Methanococcus maripaludis* Mic1c10, was provided as living culture by Amelia-Elena Rotaru (Southern University of Denmark). This strain was cultured in DSMZ medium 141 (recommended for *Methanococcus maripaludis*, but without Na_2_S) and stored at – 80 °C. Both *Methanococcus maripaludis* Mic1c10 and *Methanobacterium* strain IM1 are known corrosive methanogens, as previously documented [20, 21, 23, 24].

To start experiments, frozen stocks were revived in their respective DSMZ medium and transferred to the same medium once. When the late exponential growth phase was reached, 10 mL of culture was transferred to a sterile glass tube with conical bottom, which had first been hermetically sealed using a rubber stopper and flushed with N_2_. These tubes were then centrifuged at 2500 rpm for 15 minutes. The cell pellet was washed two times using limited medium (described below), before being resuspended in the same limited medium. For the H_2_ threshold experiment, this cell suspension was diluted with limited medium until an optical density (OD) (method described below) between 0.10 and 0.15 was reached. This procedure was used both for strains that grow planktonically or as aggregates (notably *M. barkeri* and *M. mazei*), even though similar OD values in this case do not correspond to comparable biomass densities.

All experiments were performed in limited medium. This limited medium was derived from the DSMZ medium 141 and included 0.34 g of KCl, 3.45 g of MgSO_4_ x 7 H_2_O, 4 g of MgCl_2_, 0.25 g of NH_4_Cl, 0.14 g of CaCl_2_, 0.14 g of K_2_HPO_4_, 0.2 mg of Fe(NH_4_)_2_(SO_4_)_2_, 0.2 g yeast extract powder (Y1625), 10 mL of Wolin mineral solution and 10 mL of Wolin vitamins solution for a final volume of one liter. In addition, NaCl was added for a final concentration of 0.4 g per L or 18 g per L, respectively for freshwater methanogens (*M. palustre, M. formicicum*, *M. barkeri*, *M. arboriphilus* and *M. mazei*) and marine methanogens (*strain IM1*, *M. maripaludis* DSMZ 2067 and Mic1c10 strain and *M. petrolearia*) (Table 2). In contrast to regular DSMZ medium 141, this limited medium did not contain acetate and had a ten-times lower yeast extract concentration. After autoclaving this basal medium, filter-sterile stocks solutions were added for a final concentration of 5 g of NaHCO_3_ and 0.5 g.L^−1^ of L-cysteine.

### Measurement of the H2 threshold

A volume of 80 mL of the cell suspensions obtained after washing was brought into 250 mL serum bottles. For each strain, four replicates were prepared, while three additional bottles were filled with 80 mL of limited medium as abiotic controls. The H_2_ threshold experiment was initiated by flushing the headspace of the bottles with a gas mix composed of 2 % H_2_, 78 % N_2_ and 20 % CO_2_. This mix was prepared using gas flow meters (Alicat®, MCW Series, USA) with two gas tanks, one containing 80% H_2_ and 20% CO_2_ and the other containing 80% N_2_ and 20 % CO_2_. Each bottle was pressurized at 2 bars (1 bar overpressure). The bottles were incubated upright at 30 °C with horizontal orbital shaking at 100 rpm. The total pressure was measured with a digital manometer (OMEGA DPG108, UK) before each sampling event. The headspace of each bottle was frequently sampled with a syringe for H_2_ analysis (1.5 mL). The H_2_ threshold was considered reached when the H_2_ partial pressure did not significantly decrease for at least four days with measurement intervals of maximally two days. The value of the H_2_ threshold was calculated from the average of two to four measurements performed during those last four days. After the threshold was reached, the headspace was flushed and pressurized again with the gas mix as described above. The headspace was sampled again until the H_2_ threshold was reached for a second time. The OD and the methane content in the headspace were measured at the end of the experiment. For all methanogens, the amount of CH_4_ at the end of the H_2_ threshold experiment was too low to be accurately quantified, even though detectable by GC.

### Growth yield measurements

For each methanogen, the inoculum was pre-grown and washed as described above. Four replicated bottles (250 mL) with 80 mL of limited medium were inoculated by injecting 0.5 mL of washed culture. These bottles were pressurized at 2 bars (overpressure of 1 bar) using a gas mix with 80% H_2_ and 20 % CO_2_. The bottles were incubated upright at 30 °C under horizontal orbital shaking at 100 rpm. The pressure was measured daily until at least one of the four replicates reached a negative pressure, after which the incubation was ended for all replicates. Before biomass was harvested, the headspace composition was analyzed. Next, the bottles were opened and 10 mL of each culture was transferred into a 20 mL glass tube with conical bottom and centrifuged at 3000 rpm for 15 minutes. The clear supernatant was discarded and an additional 10 mL of culture was placed in the same tube, while keeping the cell pellet. This centrifugation procedure was repeated until all 80 mL of each bottle was harvested. Each of the tubes containing the stacked cell pellets were then dried at 100 °C for 48 hours, cooled down for one hour in a desiccator and the dry weight was measured using an analytical balance (± 0.0001 g precision) (VWR, LD17 4XN). The growth yield was calculated by dividing the dried cell weight by the quantity of methane measured in the headspace at the end of the experiment.

The effect of the medium composition on the growth yield was further investigated by measuring the growth yield of one strain (*M. palustre*) in limited medium (described above) and medium with additional yeast extract and sodium acetate (final concentration of 1 g·L^−1^ each).

### Analytical methods

Gaseous samples were analyzed for H_2_ and CH_4_ using a Compact GC 4.0 (Interscience, Netherlands) with two different columns: one Molsieve 5A 30 m x 0.32 mm and one Rt-QBond 3 m x 0.32 mm, using helium as gas carrier and a TCD detector [18]. Calibration was performed using calibrating gases mixed with N_2_. The quantification limit was 1 Pa or 10 ppm for H_2_ and10 000 ppm for CH_4_. For H_2_ partial pressures below 50 ppm, a Peak Performer 1 (PP1) (Peak Laboratories, USA) GC unit based on HgO–Hg conversion with a reducing compound photometer (RCP) was also used. This GC measures trace H_2_ levels (0.1–50 ppm of H_2_) and has quantification limit well below that of the CompactGC. Partial pressures were calculated by multiplying the mol fraction, as measured by GC, with the total pressure. Dissolved concentrations were calculated from the partial pressures using the Henry constants for H_2_ at 30°C (7.8 × 10^−4^ mol.l^−1^.bar^−1^) [25]. Optical densities (OD) were measured at 600 nm using a Denovix DS-11+ spectrophotometer.

### Calculations

#### Estimation of the ATP gain from the H_2_ threshold

As explained previously [18], H_2_ thresholds can be used to calculate an estimate for the ATP gain through calculating the critical Gibbs free energy, i.e. the Gibbs free energy change when the H_2_ threshold is reached [1].

Considering hydrogenotrophic methanogenesis:

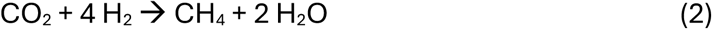

The critical Gibbs free energy (*ΔG*_c_) becomes:

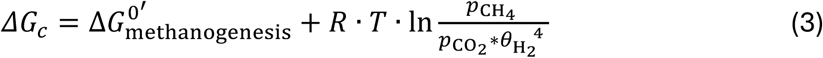

With 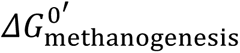 the Gibbs free energy of methanogenesis (Reaction 2) in physiological standard conditions (-131 kJ.mol^−1^ [1]), 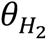 the H_2_ threshold (in atm), 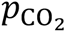 the partial pressure of CO_2_ (in atm) and 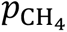 the partial pressure of methane (in atm) at the H_2_ threshold. As the methane partial pressures in the H_2_ threshold experiment were too low to be quantified, they were calculated from the amount of H_2_ consumed, according to the stoichiometric ratios of Reaction 2.

The critical Gibbs free energy at the H_2_ threshold is not zero, because methanogenesis should be considered as coupled to ADP to ATP phosphorylation (Reaction 1). At the H_2_ threshold, it can be assumed that the total Gibbs free energy change of Reaction 1 considering both methanogenesis and ADP phosphorylation, equals zero [1]:

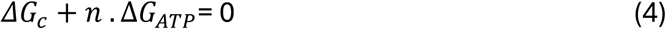

Where *ΔG*_*ATP*_ is the Gibbs free energy change for the phosphorylation of ADP into ATP. The value of this *ΔG*_*ATP*_ can vary between microorganisms and different values are used in literature [1, 18], but here we used the value of −75 kJ.mol^−1^ [26]. The ATP gain *n, i.e.* the number of ADP molecules phosphorylated for each completed methanogenesis reaction, can finally be calculated using:

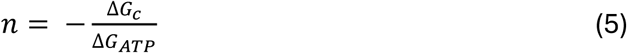

#### Estimation the ATP gain from the growth yield

The ATP gain *n* can also be estimated from experimental growth yields (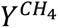, g dry cell weight (DCW) per mol methane), using the biomass yield per mol ATP (*Y*^*ATP*^, g dry cell weight per mol ATP):

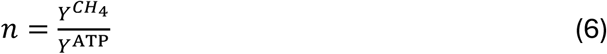

Different *Y^ATP^* values reported for methanogenesis are summarized in Table S3. Most of these values are maximum values, meaning that the actual Y_ATP_ must be lower. As our experimental growth yields are observed (net) growth yields and not maximum growth yields (requiring correction for maintenance energy), we selected a value lower than the reported maximum and used 5 g dry cell weight per mole of ATP as *Y*^ATP^ [27].

## Statistical Analysis

### Welch ANOVA test

Welch-Anova tests were performed to differentiate statistically significant groups using both GraphPad Prism® and RStudio (ver. 4.4.3). The Welch test was chosen because the variances were considered too different to perform a regular ANOVA test. All the tests were performed with α = 0.05 and the p-value was set at 0.05 for all tests.

### Error propagation

Standard deviations on the experimental results were calculated using Excel and were further propagated through the different calculations using available calculators [28, 29].

## Results and Discussion

### H2 threshold measurements

In this study, the H_2_ threshold of nine methanogenic strains was measured, as the H_2_ partial pressure at which their H_2_ consumption halts. In all abiotic controls, the H_2_ partial pressure remained almost stable for the entire duration of the experiment (Figure 2, Figure S1), as only a small drop of the total pressure was observed, as the result of headspace sampling (Figure S1).

**Figure 2:**
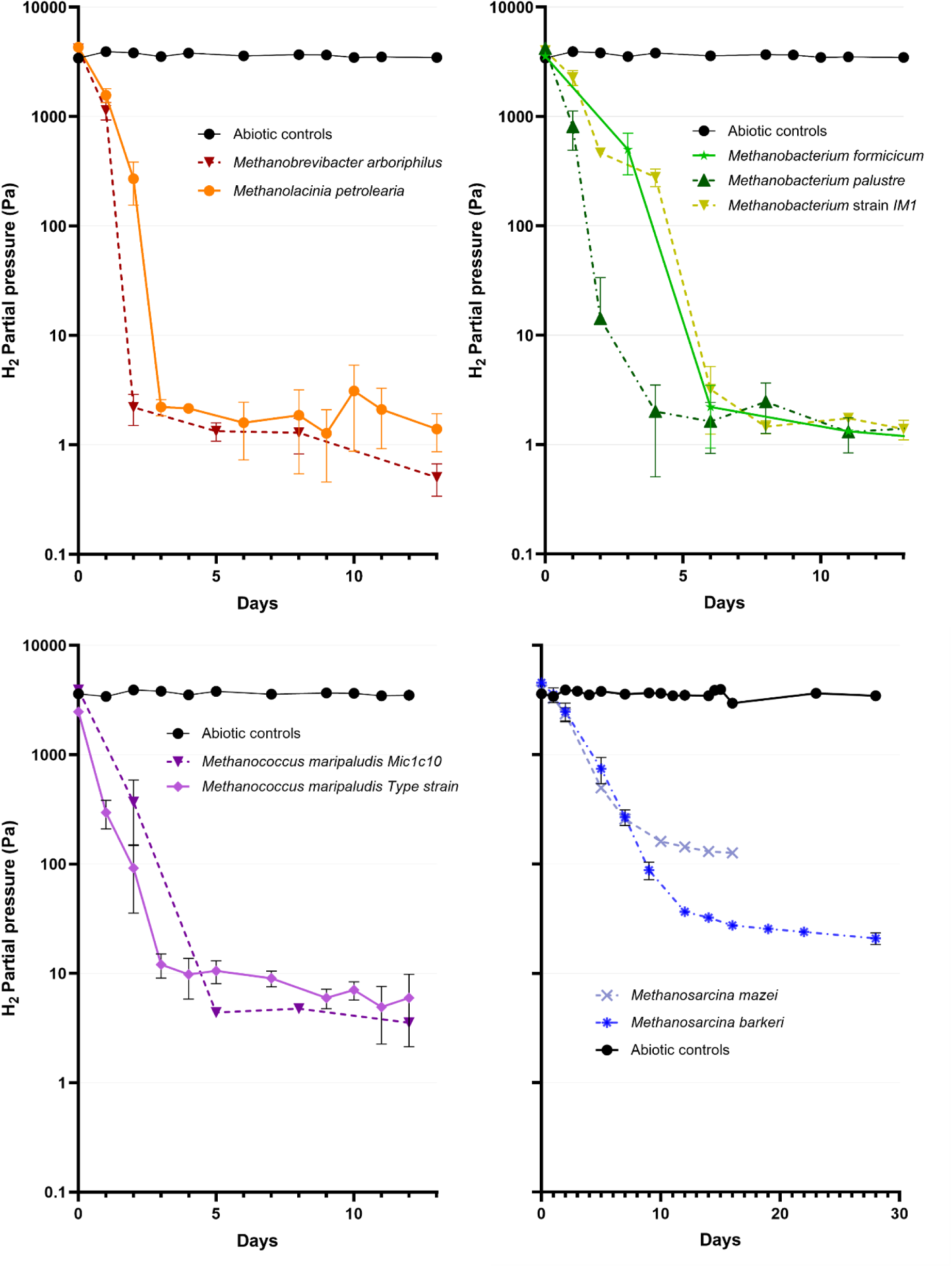
Change of the H_2_ partial pressure (Pa) over time in the H_2_ threshold experiment for the nine methanogenic species and compared to the abiotic controls. The data points represent the average of four replicates; the error bars show the standard deviations. Only datapoints before the re-addition of H_2_ are shown, later datapoints are shown in Figures S1 and S2.

For all inoculated bottles, a clear drop in the H_2_ partial pressure was observed (Figure 2), confirming H_2_ consumption by the different methanogens. Moreover, H_2_ consumption halted at a certain H_2_ partial pressure for each strain. To confirm these H_2_ thresholds, H_2_ was reinjected, once stable H_2_ partial pressures were reached. For most methanogenic strains, the H_2_ thresholds reached after the re-addition of H_2_ were comparable to those before the re-addition, except for *Methanosarcina mazei* and *Methanosarcina barkeri* (Figure S2). *M. mazei* consumed H_2_ after the H_2_ re-addition, but its H_2_ consumption halted at a higher H_2_ partial pressure (260 ± 49 Pa versus 120 ± 10 Pa before the re-addition). In contrast, *M. barkeri* no longer consumed H_2_ after the re-addition. Possibly, this is due to the loss of activity of these strains over the long incubation times (*M. mazei* reached its H_2_ threshold after 12 days, while *M. barkeri* required 28 days). All other methanogens required only five days or less to reach their H_2_ threshold.

The three *Methanobacterium* species displayed similar low H_2_ thresholds around 2 Pa (Figure 2, Figure 3, Table 2). Also, *Methanosarcina petrolearia* and *Methanobrevibacter arboriphilus* reached H_2_ thresholds in this range. *M. arboriphilus* was the only species that reached a H_2_ partial pressure of 1 Pa (Figure 2, Figure 3, Table 2). The Welch-ANOVA test gathered these five strains with the lowest H_2_ thresholds in one statistical group (Figure 3). In contrast, all the other strains had higher H_2_ thresholds and were classified each as a significantly different group (Figure 3). The two strains of *Methanococcus maripaludis* displayed H_2_ thresholds slightly higher than the five strains with the lowest H_2_ thresholds (Figure 2, Figure 3, Table 2). Interestingly, the corrosive strain *Mic1c10* had a significantly lower H_2_ threshold (around 4 ± 1 Pa) than the type strain (7 ± 1 Pa).

**Figure 3:**
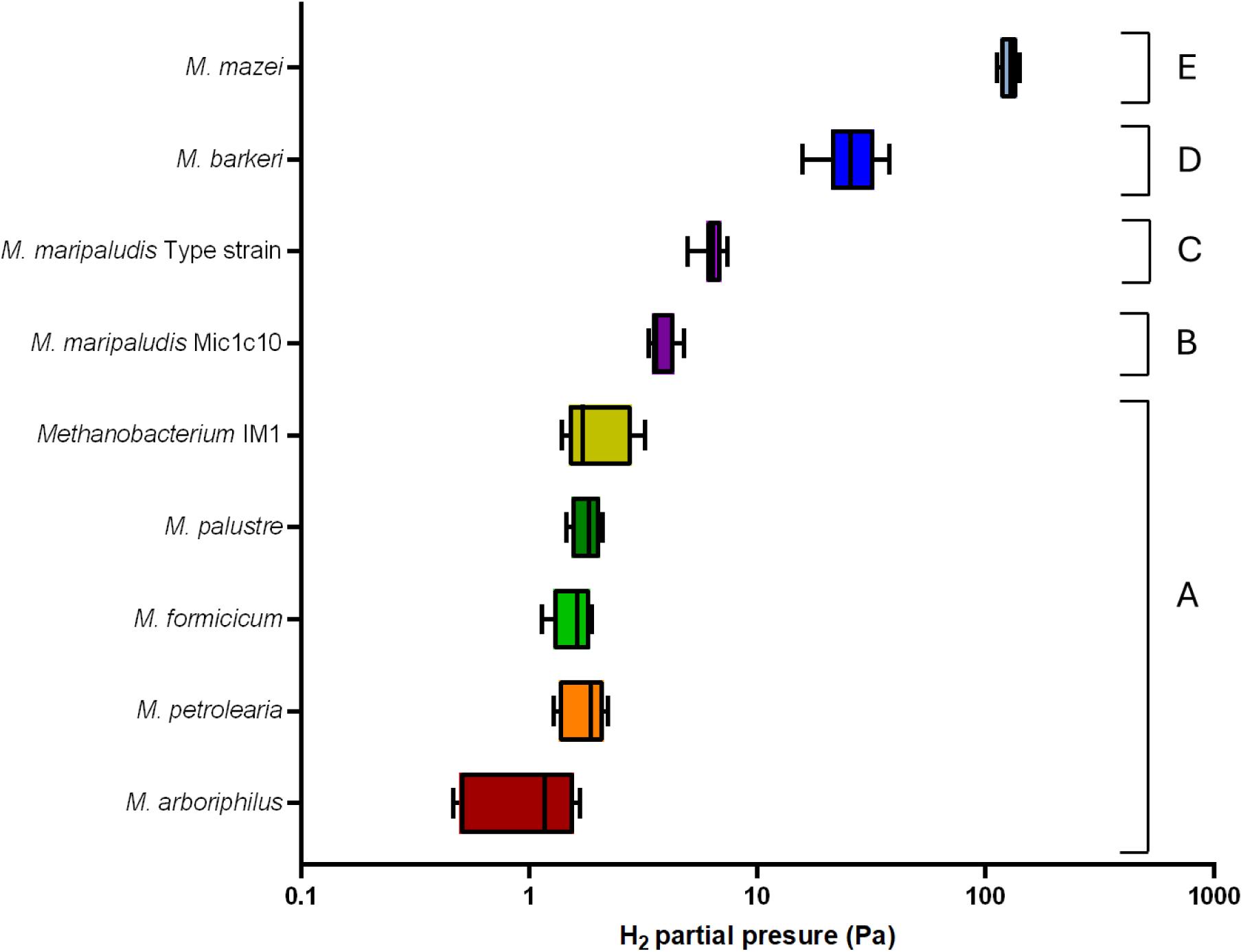
Box plots summarizing the experimental H_2_ threshold values for the nine methanogenic strains. The letters indicate statistically different groups, as determined using a Welch-ANOVA test.

The *Methanosarcina* species differed strongly in their H_2_ thresholds, as *M. barkeri* had a H_2_ threshold of 21 Pa ± 2 Pa, while the highest H_2_ threshold was recorded for *M. mazei* (120 ± 10 Pa). Previously, widely varying H_2_ thresholds were reported for *M. barkeri* (Table S1). These variations are possibly due to different experimental conditions (Table S1) or the slow H_2_ consumption by this strain. Our H_2_ threshold for *M. barkeri* could be somewhat overestimated, as this strain could not be reactivated after the halt of H_2_ consumption. In contrast, *M. mazei* was still active after the H_2_ re-addition, so its six-times higher H_2_ threshold in comparison to *M. barkeri* (Table 2) cannot be due to an experimental artefact.

Overall, we obtained experimental H_2_ thresholds ranging over two orders of magnitude (Figure 3, Table 2), which is a similar range as previously recorded (Table S1). The H_2_ thresholds also agree with the classification based on the presence of cytochromes (Table 1). We found H_2_ thresholds below 7 Pa for all methanogens without cytochromes, while methanogens with cytochromes had H_2_ thresholds above 20 Pa (Table 2). Our results thus differentiate both groups more strongly than previous recordings (Table 1). Moreover, in both groups, we found methanogens with statistically different H_2_ thresholds (Figure 3), showing that additional variations in H_2_ thresholds exist.

### Growth yield measurements

In parallel of the H_2_ threshold study, all methanogenic strains were grown in bottles on 80% H_2_ and 20% CO_2_ to determine their growth yields. In all these bottles methane levels significantly increased over the incubation period, while H_2_ was consumed and cell growth occurred (Table S4). These results were used to calculate the growth yield, i.e. the increase of biomass expressed as gram dry cell weight (g_DCW_) versus the amount of methane produced (Figure 4, Table 2). For one strain, *M. palustre*, the growth yield was also determined with additional acetate and yeast extract in the medium (1 g·L^−1^ each). A significantly higher growth yield was found with the addition of extra organic carbon sources (Figure S3), demonstrating the importance of using the same medium conditions when comparing growth yields.

**Figure 4:**
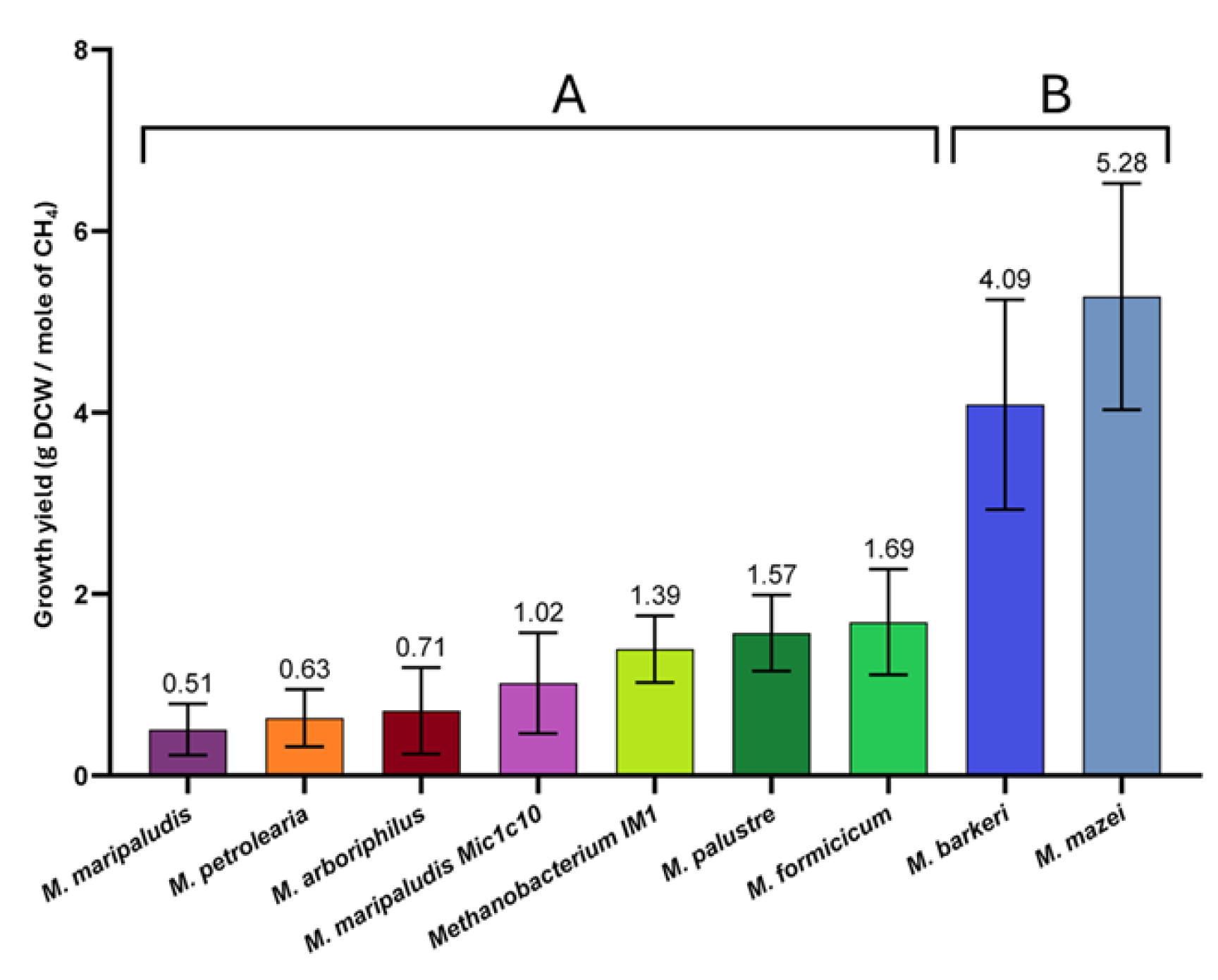
Comparison of the growth yields expressed as the increase of gram Dry Cell Weight (g_DCW_) per mole of methane produced for the different methanogenic strains. The letters indicate statistically different groups, as determined using a Welch-ANOVA test.

Relatively high variabilities between the replicates were obtained for the growth yields (large error bars in Figure 4). This is likely due to the low total weight of dried biomass (in mg) obtained during the growth yield experiment (Table S4), limiting the accuracy of this measurement. In addition, the centrifugation steps were possibly not always as effective in removing cells from the culture suspension, which could also explain the high variability between the replicates and could have underestimated some of the growth yields. A high variability is fairly common for growth yield measurements, as this was also observed in other studies determining methanogenic growth yields [11, 30, 31].

Despite the high variability in the dry cell weight, Welch-ANOVA tests identified two significantly different groups in the growth yields (Figure 4, right). The two *Methanosarcina* strains form one group with high growth yields of 4.09 ± 1.16 and 5.28 ± 1.25 g_DCW_/mole of CH_4_ for *M. barkeri* and *M. mazei* respectively. All other methanogens had significantly lower growth yields, ranging from 0.51 ± 0.28 g_DCW_/mole of CH_4_ (for *M. maripaludis* type strain) to 1.69 ± 0.58 g_DCW_/mole of CH_4_ (for *M. formicicum*). Growth yields were also expressed as g_DCW_ per mole H_2_ consumed (Figure S4), leading to the same statistical groups.

Overall, our growth yields are consistent, but in the lower range, of previously reported growth yields (Table 2 and S2), likely because our medium did not contain organic carbon sources (Figure S3). Moreover, the two significantly different groups identified in the growth yields agree with the classification based on the presence of cytochromes, while our results differentiate both groups more strongly than previous findings (Table 1). In both groups, no further significant differences in growth yields were identified. Even for *M. mazei* and *M. barkeri*, which differ six times in their H_2_ threshold, no significant different growth yields were found (Figure 4). It is likely that this incoherence is due to the high variability of the growth yield measurement (high error bars in Figure 4).

### Comparison of ATP gain estimates

Both the H_2_ threshold and the growth yield allow to estimate the ATP gain. Using the H_2_ threshold, the ATP gain estimates ranged from 0.32 ± 0.07 for *M. arboriphilus* to 0.97 ± 0.01 for *M. mazei* (Table 2). Estimating ATP gains from the growth yields gave a slightly wider range, starting at 0.10 ± 0.03 for *M. maripaludis* up to 1.06 ± 0.12 for *M. mazei*. Despite the totally different experimental procedures and the various assumptions underlying these calculations (Equations 5 and 6), both estimates led to a largely overlapping range, validating our approach. These ATP gain estimates were further correlated to each other, leading to a clear but not perfect correlation (Figure 5, R^2^ of 0.84). Only the two *Methanosarcina* species, *M. mazei* and *M. barkeri,* fall on the 1:1 correlation line. For all other strains, higher ATP gains were calculated from the H_2_ thresholds than from the growth yield, demonstrating that there were discrepancies between our methods, possibly because the growth yields were somewhat underestimated. This illustrates that our ATP gains should be considered as estimates, rather than absolute values. Nevertheless, the clear correlation (Figure 5) does suggest that the ATP gains differed among the tested strains.

**Figure 5:**
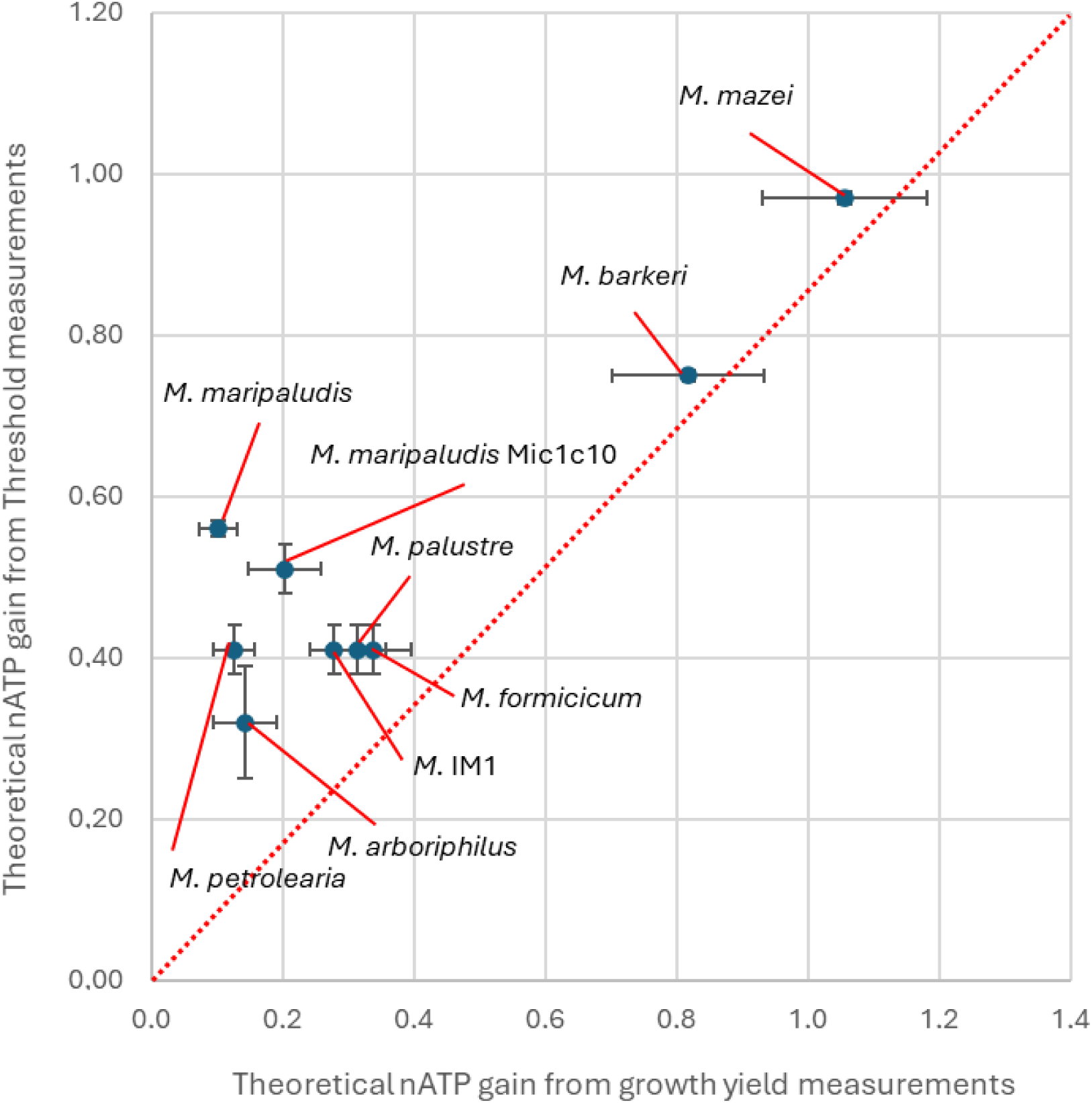
Correlation of the ATP gains estimations from the experimental H_2_ thresholds versus the ATP gains estimated from the experimental growth yields. The vertical and horizontal error bars reflect the standard deviation on these estimates and were propagated from the experimental errors. The red line indicates a 1:1 correlation. The correlation coefficient (R2) was 0.84.

These ATP gain estimates also agree with the cytochrome-dependent classification of methanogens (Table 1 and 2). Kaster et al. [7] estimated an ATP gain higher than 1 for *M. barkeri*, while we calculated this to be 0.82 ± 0.11 ATP per mol CH_4_, when derived from the growth yield (Table 2). For methanogens without cytochromes, Kaster et al. [7] proposed an ATP gain lower than 1, while Goyal et al. [32] estimated an ATP gain of 0.35 for *M. maripaludis*. Both correspond well with the range of ATP gain estimates (between 0.10 and 0.56 mol ATP per mol CH_4_) determined here for methanogens without cytochromes (Table 1 and 2).

In addition, our ATP gain estimates calculated from the H_2_ thresholds suggest that additional variations in energy conservation exist, as additional significant differences in H_2_ thresholds were recorded in both groups of methanogens (Figure 3). Unfortunately, this was not confirmed by the ATP gains derived from the growth yields, likely again due to the high variability on the growth yields. Besides the two different mechanisms for energy conservation in methanogens (Figure 1), various additional variations in energy conservation components are currently being unraveled [6, 8, 9, 33]. Moreover, *Methanothermobacter marburgensis* was recently shown to have a flexible energy conservation system, adapting to the nickel concentration in the medium [34]. We therefore conclude that the measurement of H_2_ thresholds could offer a valuable tool to assess how variations in energy conservation affect the ATP gain.

### Link between the H2 threshold and electroactivity of methanogens

The experimental H_2_ thresholds obtained in this study allow to reevaluate previously reported differences in electroactivity among methanogens. Mayer et al [19] studied the electromethanogenesis performance of several methanogens. *M. maripaludis* and *M. petrolearia*, for which a low H_2_ threshold was found here (Figure 3), increased cathodic currents (i.e. key signature of electroactivity), just as several other methanogens without cytochromes. In contrast, *M. mazei*, for which the highest H_2_ threshold was recorded in this study (Figure 3), was incapable of electromethanogenesis [19]. Similarly, Jaramillo *et al*. investigate the corrosion potential of different methanogens [35]. Non-hydrogenotrophic methanogens were reported as non-corrosive, confirming the important role of H_2_ as intermediate in microbial corrosion. *M. mazei* and *M. barkeri* (high H_2_ threshold) were mild corrosive, whereas *Methanobacterium* spp. (low H_2_ threshold) were highly corrosive. Overall, these correlations thus suggest that methanogens with a low H_2_ threshold are more electroactive than methanogens with a high H_2_ threshold.

Nevertheless, several other variables likely affect the electroactivity of methanogens, in addition to the H_2_ threshold. The H_2_ pressure maintained at the cathode or steel surface also depends on the rate of H_2_ consumption. Moreover, the H_2_ evolution reaction on a cathode or steel surface is stimulated by extracellular enzymes, such as hydrogenases. *M. maripaludis* Mic1c10 has a unique extracellular hydrogenase anchored to its cell surface [23], causing stronger corrosion than other *M. maripaludis* strains [23]. Moreover, its cathodic current increase was double as high than for a non-corrosive *M. maripaludis* strain [19, 36]. Here, we found that strain Mic1c10 had a significantly lower H_2_ threshold than the *M. maripaludis* wild type (Figure 3, Table 2). Kawaichi et al [23] also recorded that strain Mic1c10 maintained slightly lower H_2_ levels during steel corrosion than by the *M. maripaludis* type strain. These differences, however, are possibly too small to explain the strong corrosion rates caused by Mic1c10, which is thus likely mostly due to its extracellular hydrogenases. Similarly, *Methanobacterium* -IM1 was more corrosive than other tested methanogens [24], but here we did not find that strain IM1 had a lower H_2_ threshold than other tested *Methanobacterium* strains (Figure 2, Table 3). *Methanobacterium* IM1 thus likely has an additional mechanism to enhance extracellular electron uptake rate, which is possibly similar to the one described for *M. maripaludis* Mic1c10.

Overall, we conclude that the electroactivity of methanogens is inversely proportional to their H_2_ threshold, whereas few methanogens (e.g. *M. maripaludis* Mic1c10, *Methanobacterium* IM1) have additional mechanisms to increase their electroactivity.

### Ecological and biotechnological implications of variations among methanogens

We systematically found that methanogens with a low H_2_ threshold have a low growth yield and a low ATP gain (Table 2). In contrast, methanogens with a high H_2_ threshold have a high growth yield and have a higher ATP gain (Table 2). Hydrogenotrophic methanogens are thus confronted with a trade-off between the capacity to consume H_2_ at trace levels or the ability to grow strongly (high growth yield) with a high energetic efficiency (high ATP gain). This trade-off allows to understand the ecological niche and biotechnological relevance of the different types of methanogens. For instance, Mills *et al.* [37] found that the presence of the *Methanosarcina* genus in electro-methanogenesis systems was correlated to high H_2_ levels (reactors with low coulombic efficiency), while the *Methanobacterium* genus was rather correlated to low H_2_ levels (high coulombic efficiency).

In addition, a correlation between the growth yield (high ATP gain) and the growth rate of methanogens would be expected [18]. However, Thauer [1] described that methanogens with cytochromes (high ATP gain) have a lower growth rate (high doubling time) than methanogens without cytochromes (low ATP gain, but low doubling times). Comparing doubling times is though not adequate, since methanogens strongly vary in cell size (large clustered cells for *Methanosarcina* species versus small different shaped cells for other methanogens) [38, 39]. Nevertheless, also in the H_2_ threshold experiment of this study, H_2_ was faster consumed by methanogens without cytochromes, than by methanogens with cytochromes (Figure 2). This observation possibly reflects that *Methanosarcina* species are not well adapted to consume H_2_ at lower levels. These methanogens are also capable of acetoclastic and methylotrophic methanogenesis, in contrast to the other tested methanogens. Methanogens with cytochromes are thus possibly mainly adapted to use other substrates than H_2_ and only shift to H_2_ consumption at high H_2_ partial pressures.

In this study, we found that H_2_ thresholds of methanogens range over two orders of magnitude (Figure 3). In comparison, we previously found three orders of magnitude difference in the H_2_ thresholds of acetogenic bacteria (6 Pa to 1990 Pa) [18]. It is often assumed that acetogens cannot compete with methanogens for low H_2_ levels, as the latter have lower H_2_ thresholds [40]. Our results, however, demonstrate that such a competition rather should be evaluated on species level. For instance, *Methanobacterium* strains indeed have lower H_2_ thresholds than *Acetobacterium* spp, but acetogenic *Sporomusa* spp. have H_2_ thresholds in the range of those of methanogens without cytochromes and well below those with cytochromes. The outcome of competition for trace H_2_ levels between methanogens and acetogens likely also depends on the H_2_ consumption rates. Recently, H_2_ consumption by acetogens was described by first order kinetics (increasing rates at increasing H_2_ levels) [41], while the kinetics of H_2_ consumption by methanogens rather follows Monod kinetics with low half-saturation constants [42] or even zero-order kinetics [43]. Based on these kinetic differences, methanogens (without cytochromes) consume H_2_ likely faster at low H_2_ levels, while acetogens have a faster H_2_ consumption at higher H_2_ levels, even though H_2_ consumption rates can still widely vary within these groups [41].

Overall, we conclude that the trade-off between a low H_2_ threshold and high growth yield of hydrogenotrophic methanogens explains their ecological prevalence and affects their biotechnological applicability.

## Supporting information

Supplementary Informations

## Acknowledgement

This study was financed by a Villum Young Investigator grant (VIL53083). We would like to thank Satoshi Kawaichi and Amelia-Elena Rotaru from the University of Southern Denmark for providing the living culture of *Methanococcus maripaludis* Mic1cc10 and Florin Musat from Aarhus University for providing the living culture of *Methanobacterium* IM1. We also acknowledge the Section for Microbiology at Aarhus University for lending their Peak Performer. Heidi Skov Johansen and Laura Mũnoz-Duarte are thanked for their practical support during the experiments and calculations. We also appreciate the advice of Sanne Sandberg Overby on the statistical tests.

## Data availability statement

The data that supports the findings of this study are available in the supporting information file.

## Conflict of interest statement

The authors declare no conflict of interest.

## Notes

### Competing Interest Statement

The authors have declared no competing interest.

### Summary of Updates

The experimental results remain the same but some of the interpretations have been removed, notably the Carbon assimilation part in the discussion.

