## Supplementary Informations for "Variations in H_2_ thresholds and growth yields reveal bioenergetic diversity among hydrogenotrophic methanogens"

**Supplementary information**

**to**

### Variations in $H_2$ thresholds and growth yields among hydrogenotrophic methanogens relate to different energy conservation strategies

Timothé Philippon [(1)]. Jo Philips [(1). (2)] *

[(1)] Department of Biological and Chemical Engineering. University of Aarhus.

Gustav Wieds Vej 10. 8000 Aarhus C. Denmark

[(2)] Novo Nordisk Foundation $CO_2$ Research Center (CORC). Gustav Wieds Vej 10.

[(3)] 8000 Aarhus C. Denmark

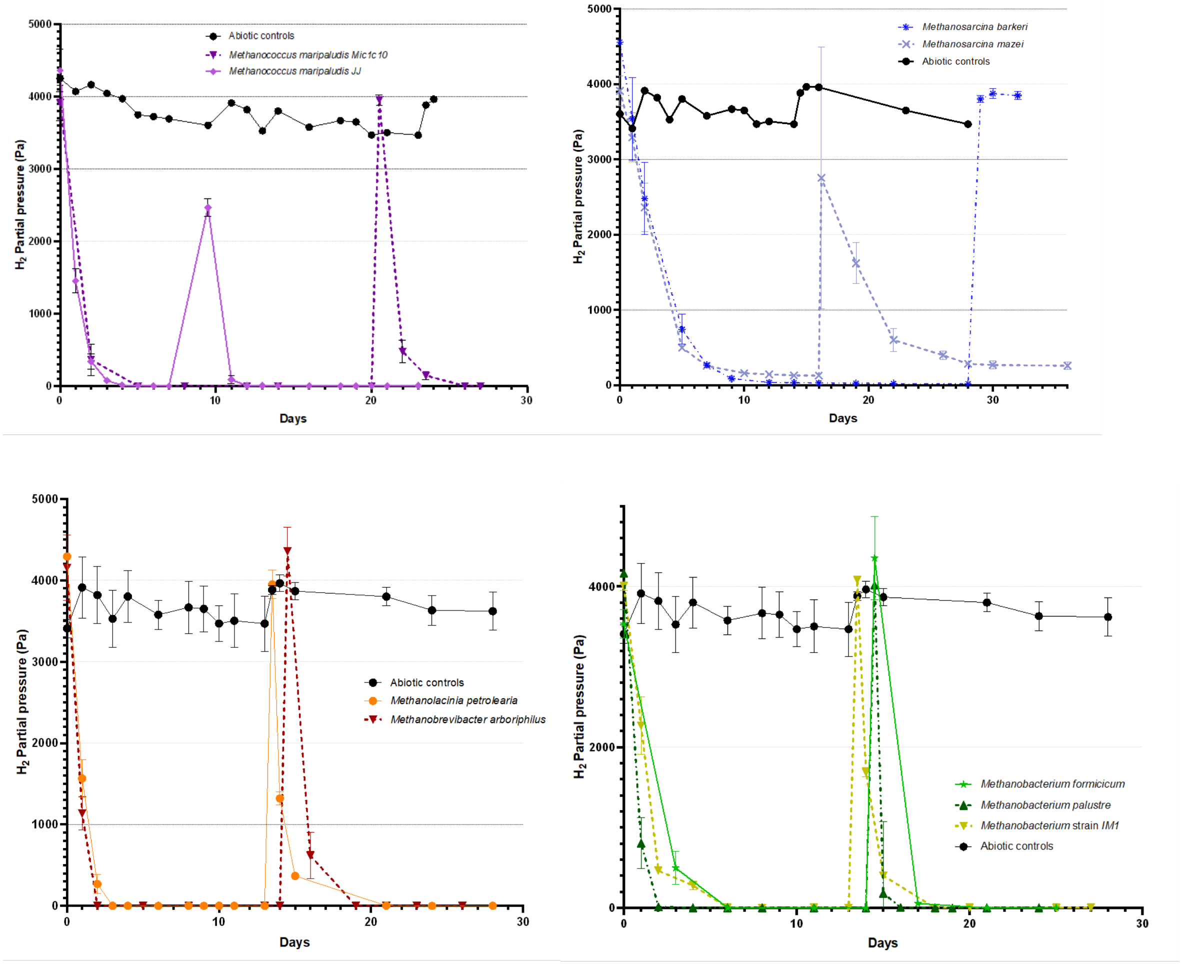


Figure S1: Change of the $H_2$ partial pressure (Pa) over time in the $H_2$ threshold experiment in linear scale for the nine methanogenic species and compared to the abiotic controls including the refill of the headspace; the error bars show the standard deviations.

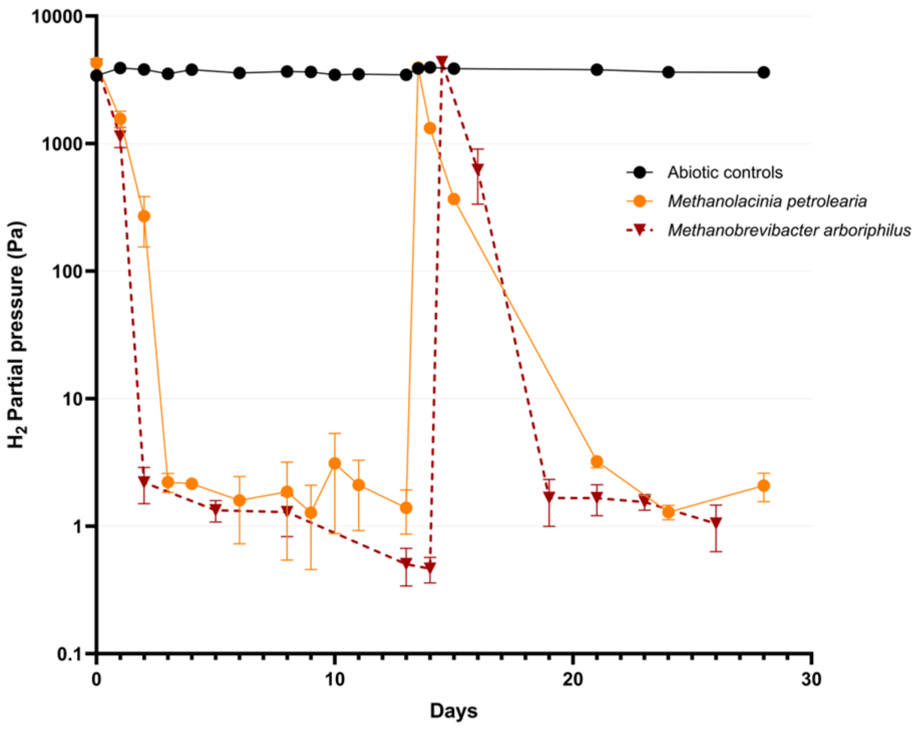

10000
1000
100
10
1
0.1
0
10
20
30
Days
$H_2$ Partial pressure (Pa)
Abiotic controls
*Methanolacinia petrolearia*
*Methanobrevibacter arboriphilus*


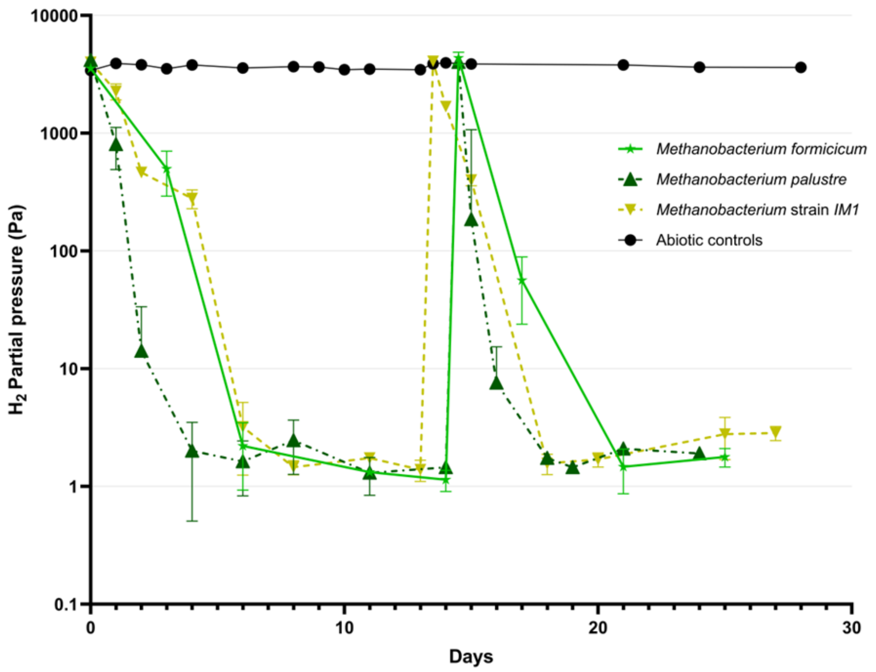

10000
1000
100
10
1
0.1
0
10
20
30
Days
$H_2$ Partial pressure (Pa)
*Methanobacterium formicicum*
*Methanobacterium palustre*
*Methanobacterium* strain *IM1*
Abiotic controls

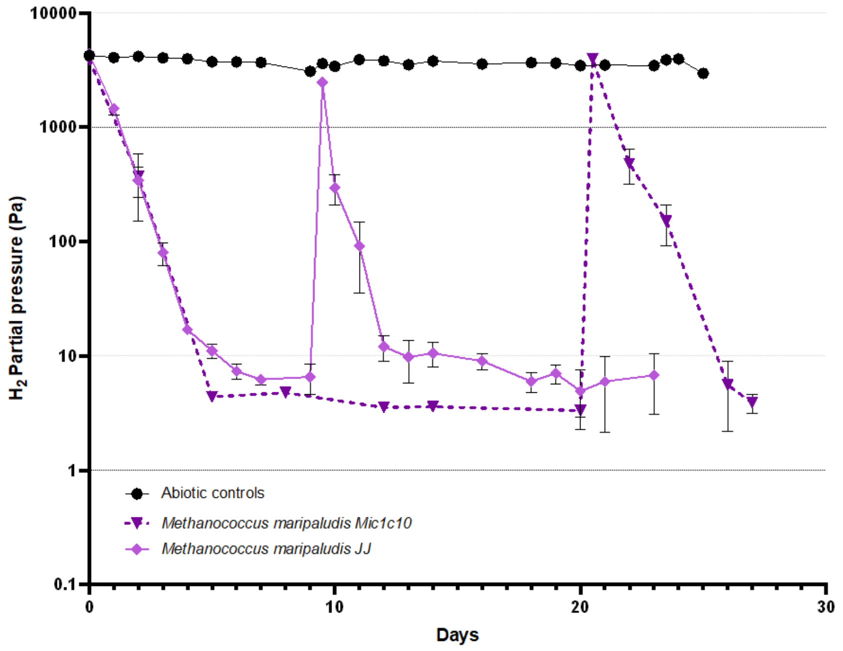


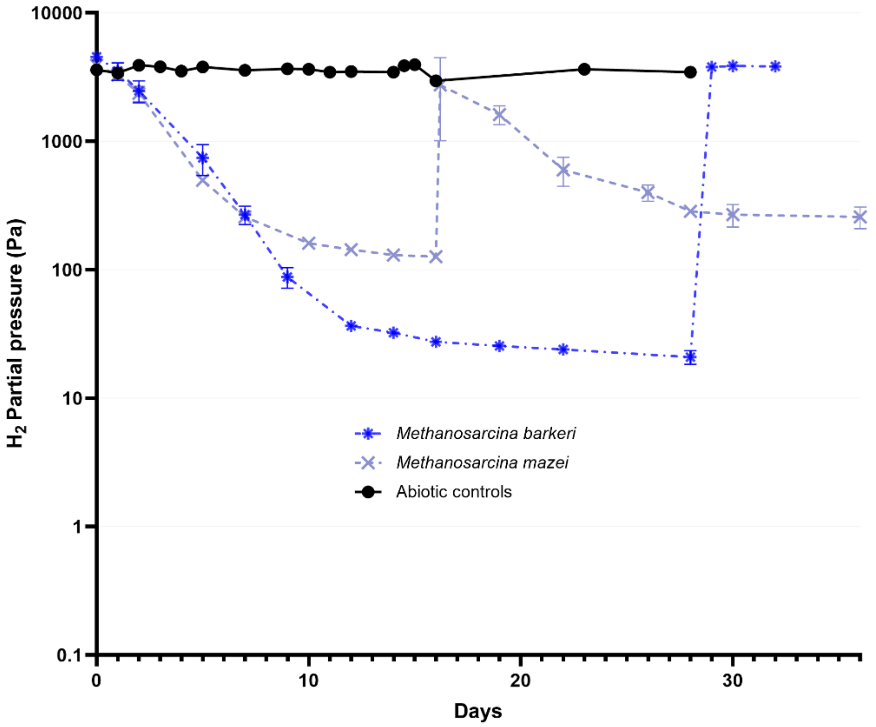


Figure S2: Change of the $H_2$ partial pressure (Pa) over time in the $H_2$ threshold experiment for the nine methanogenic species and compared to the abiotic controls including the refill of the headspace; the error bars show the standard deviations. Note the logarithmic scale of the Y-axis.

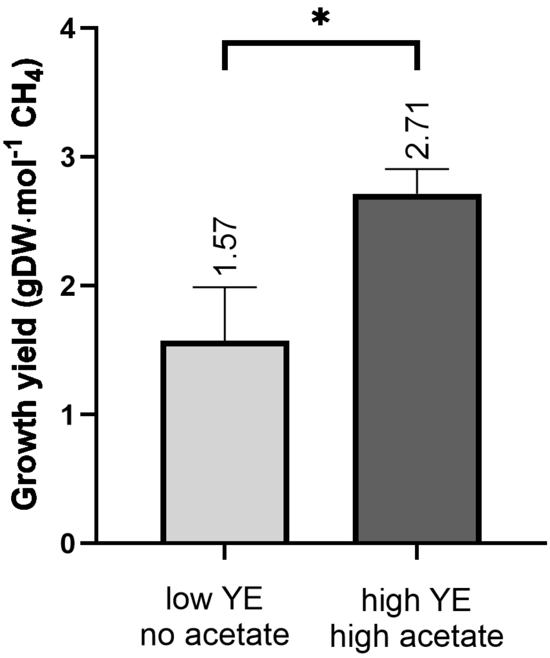


Figure S3: Comparison of the growth yield of M¡.palustre expressed as amount of biomass (in $g_{DCW}$) per mole of $CH_4$ produced, when grown either (left) in limited medium containing 0.2 $g{\cdot}L^{-1}$ yeast extract (YE) and no sodium acetate; or (right) in complex medium containing 1.0 $g{\cdot}L^{-1}$ YE and 1.0 $g{\cdot}L^{-1}$ sodium acetate. The star indicates a statistically significant difference between the two groups, as determined using a student t-test ($P < 0.01$).

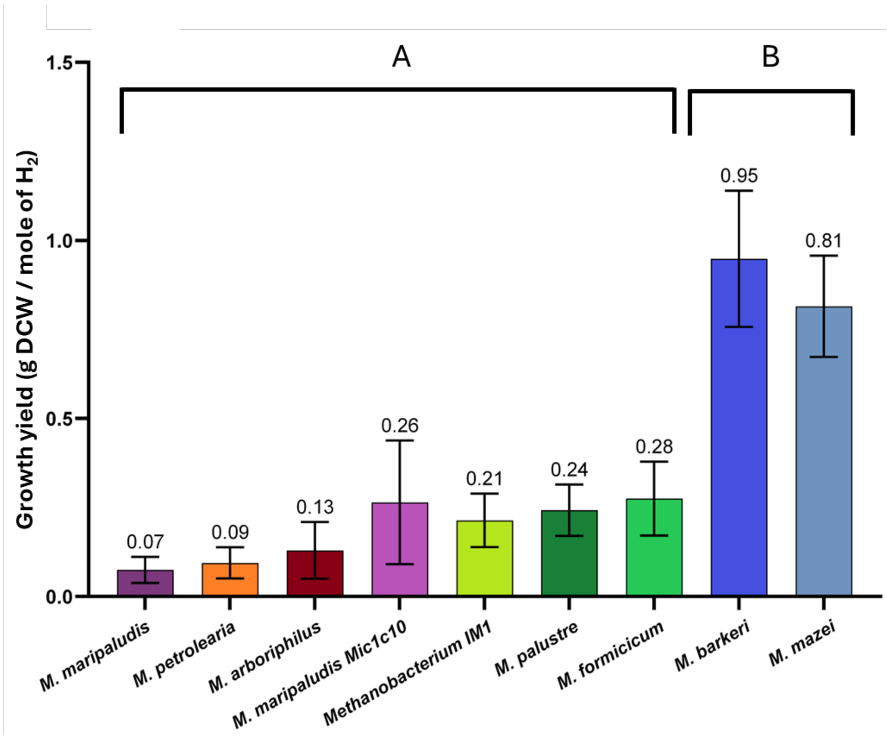


Figure S4: Comparison of the growth yield for all the methanogenic species expressed as the amount of biomass (in $g_{DCW}$) per mole of $H_2$ consumed. The letters indicate statistically different groups, as determined using a Welch-ANOVA test.

Table S1: Overview of previously reported $H_2$ thresholds for hydrogenotrophic methanogens, including experimental conditions. These studies used different initial $H_2$ concentrations, resulting in varying methane partial pressures and total pressures when the $H_2$ threshold was reached.

| Methanogen | $H_2$ threshold (Pa) | Conditions | References |
|---|---|---|---|
| Methanogens without cytochromes | | | |
| Methanobacterium formicicum | 4 .5±2.1 | 37 °C, 100% CO2, 3 atm, starting with 3000-6000 Pa $H_2$ | (Kral et al., 1998) |
| | 6.5 ± 0.6 | 39 °C, 20% $CO_2$, 2.3 atm, starting with 500-2000 Pa $H_2$ | (Lovley, 1985) |
| | 5.6 | 28-34 °C, 20% $CO_2$, 2 atm, starting with 2000 Pa $H_2$ | (Cord-Ruwisch et al., 1988) |
| Methanobacterium bryantii | 6.9 ± 1.5 | 39 °C, 20% $CO_2$, 2.3 atm, starting with 500-2000 Pa $H_2$ | (Lovley, 1985) |
| | 0.87-3.19 | 37°C, 20% $CO_2$, 0.2 atm, starting with 80% $H_2$, depending on nickel concentration | (Neubeck et al., 2016) |
| Methanobrevibacter arboriphilus | 18 | 28-34 °C, 20% $CO_2$, 2 atm, starting with 2000 Pa $H_2$ | (Cord-Ruwisch et al., 1988) |
| | 8 | 37 °C, 20% $CO_2$, 1 atm, starting with 30 Pa $H_2$ | (Kaster et al., 2011) |
| Methanobrevibacter smithii | 20 | 28-34 °C, 20% $CO_2$, 2 atm, starting with 2000 Pa $H_2$ | (Cord-Ruwisch et al., 1988) |
| Methanobrevibacter strain AMG1 | 5.7±0.7 | 30 °C, 20% $CO_2$ starting with 150 Pa $H_2$ | (Feldewert et al., 2020) |
| Methanospirillum hungatei | 9.5 ± 1.3 | 39 °C, 20% $CO_2$, 2.3 atm, starting with 500-2000 Pa $H_2$ | (Lovley, 1985) |
| | 6 | 28-34 °C, 20% $CO_2$, 2 atm, starting with 2000 Pa $H_2$ | (Cord-Ruwisch et al., 1988) |
| Methanococcus maripaludis | 9±3 | 25 °C, 100% CO2, 3 atm, starting with 3000-6000 Pa $H_2$ | (Kral et al., 1998) |
| Methanococcus vannielii | 15 | 28-34 °C, 20% $CO_2$, 2 atm, starting with 2000 Pa $H_2$ | (Cord-Ruwisch et al., 1988) |
| Methanoculleus bourgensis MAB1 | 0.10-0.17 | 37°C, 20% $CO_2$, 0.2 atm, starting with 80% $H_2$, depending on nickel concentration | (Neubeck et al., 2016) |
| Methanogens wit cytochromes | | | |
| Methanosarcina barkeri | 47±6 | 37 °C, 100% CO2, 3 atm, starting with 3000-6000 Pa $H_2$ | (Kral et al., 1998) |
| | 13-22 | 37°C, 20% $CO_2$, 0.2 atm, starting with 80% $H_2$, depending on nickel concentration | (Neubeck et al., 2016) |
| | 20-35 | 0 °C, 92.5% $CO_2$, 0.007-0.012 atm, Starting with 2.9% $H_2$ | (Harris and Schuerger, 2025) |
| | 150 | 37 °C, 20% $CO_2$, 1 atm, starting with 600 Pa $H_2$ | (Kaster et al., 2011) |

Table S2: Overview of previously reported growth yields for methanogens growing on $H_2$ and $CO_2$, including their dependence on the growth conditions. In addition to different medium compositions, these studies used various experimental setups to determine growth yields.

| Methanogen | Growth yield ($g_{DCW}\cdot mole^{-1}$ $CH_4$) | Growth conditions | Experimental setup | Reference |
|---|---|---|---|---|
| Methanogens without cytochromes | | | | |
| Methanobacterium.formicicum | 3.5 | Medium with 0.5 $g\cdot L^{-1}$ YE and 0.5 $g\cdot L^{-1}$ trypticase | Fed-batch fermenter with gas sparging | (Schauer and Ferry, 1980) |
| Methanobrevibacter.arboriphilus | 2.5 | Medium with 2 $g\cdot L^{-1}$ YE and 1 $g\cdot L^{-1}$ tryptone | Fed-batch fermenter with gas sparging | (Kaster et al., 2011) |
| | 1.06–1.42 | Mineral medium, depending on growth rate | Continuously operated chemostat | (Morii et al., 1987) |
| Methanococcus.maripaludis | 2.86 | Medium with 2 $g\cdot L^{-1}$ casamino acids | Continuously operated chemostat | (Costa et al., 2013) |
| | 0.77 | Medium with 2 $g\cdot L^{-1}$ casamino acids, but limited in ammonia and phosphate | Continuously operated chemostat | (Costa et al., 2013) |
| | 2.2-3.0 | Medium with acetate as carbon source, depending on growth rate | Continuously operated chemostat | (Richards et al., 2016) |
| | 3.54 ± 0.15 | Mineral medium | Batch bottles | (Goyal et al., 2016) |
| | 1.49 | Mineral medium | Continuously operated chemostat | (Xue et al., 2025) |
| Methanothermobacter. thermoautotrophicus | 1.50-1.65 | 45-65°C, limited medium, $H_2$ and $CO_2$ gas supply not limiting | Fed-batch fermenter with gas sparging | (Schönheit et al., 1980) |
| | 3.0 | 55°C, limited medium, $H_2$ and $CO_2$ gas supply limiting | Fed-batch fermenter with gas sparging | (Schönheit et al., 1980) |
| | 1.4-3.0 | Limited medium, depending on the dissolved $H_2$ concentration | Fed-batch fermenter with gas sparging + Continuously operated chemostat | (de Poorter et al., 2007) |
| Methanogens.with.cytochromes | | | | |
| Methanosarcina.barkeri | 5.40 ± 1.87 | Mineral medium | Batch bottles | (Ferguson and Mah, 1983) |
| | 8.38 ± 0.88 | Medium with 2 $g\cdot L^{-1}$ YE | Batch bottles | (Ferguson and Mah, 1983) |
| | 7.5 | Medium with 2 $g\cdot L^{-1}$ YE and 1 $g\cdot L^{-1}$ tryptone | Fed-batch fermenter with gas sparging | (Kaster et al., 2011) |
| | 6.4 | Mineral medium | Fed-batch fermenter with gas sparging | (Weimer and Zeikus, 1978) |

Table S3: $Y_{ATP}$ values reported for methanogenesis and other metabolisms. Many of these values are reported as maximum values, meaning that the actual $Y_{ATP}$ must be lower. As our experimental growth yields are observed (net) growth yields and not maximum growth yields (requiring correction for maintenance energy), we selected a $Y_{ATP}$ lower than the reported maximum. We used a $Y_{ATP}$ value of 5 $g_{DCW}$/mol ATP to calculate the ATP gain from the experimental growth yields (Equation 6, main text).

| $Y_{ATP}$ ($g_{DCW}$/mol ATP) | Specifications | Source |
|---|---|---|
| 4-6 | Autotrophic growth | (Spormann, 2023) |
| 10 | Heterotrophic growth | |
| Max. 6.5 | Autotrophic growth for methanogens, maximum value needs to be corrected for maintenance energy; decreases with lower growth rate | (Thauer et al., 2008) |
| Max. 8 | Methanogen growing on $CO_2$ | (Hedderich and Whitman, 2013) |
| Max. 10 | Hydrogenotrophic methanogen using acetate as carbon source | |
| 28 | Heterotroph | |
| Max. 4.75 | Autotroph using Calvin cycle | |

Table S4: Substrate consumption, methane production and biomass growth observed for the different replicates of the growth yield experiment.

| | Total $H_2$ consumed (mmole) | Total $CH_4$ produced (mmole) | Dry biomass created (mg) |
|---|---|---|---|
| Methanococcus maripaludis Mutant strain Mic1c10 | | | |
| Replicate 1 | 2.98 | 0.94 | 1.60 |
| Replicate 2 | 4.37 | 1.11 | 0.60 |
| Replicate 3 | 12.85 | 2.93 | 1.20 |
| Replicate 4 | 12.77 | 2.59 | 3.70 |
| Methanosarcina barkeri | | | |
| Replicate 1 | 12.00 | 2.13 | 11.10 |
| Replicate 2 | 3.09 | 0.91 | 2.00 |
| Replicate 3 | 4.77 | 1.22 | 5.10 |
| Replicate 4 | 5.13 | 1.24 | 5.90 |
| Methanococcus maripaludis Type strain | | | |
| Replicate 1 | 13.08 | 2.20 | 1.30 |
| Replicate 2 | 13.15 | 1.76 | 1.6 |
| Replicate 3 | 11.23 | 3.06 | 0.4 |
| Replicate 4 | 11.61 | 1.25 | 0.5 |
| Methanolacinia petrolearia | | | |
| Replicate 1 | 9.67 | 1.49 | 0.6 |
| Replicate 2 | 10.80 | 1.67 | 0.7 |
| Replicate 3 | 10.23 | 1.48 | 1.6 |
| Methanobacterium IM1 | | | |
| Replicate 1 | 11.14 | 2.00 | 3.8 |
| Replicate 2 | 11.96 | 2.10 | 1.8 |
| Replicate 3 | 8.87 | 1.05 | 1.5 |
| Replicate 4 | 11.81 | 1.65 | 2.3 |
| Methanobrevibacter arboriphilus | | | |
| Replicate 2 | 12.82 | 2.20 | 2.9 |
| Replicate 3 | 13.71 | 2.71 | 1.8 |
| Replicate 4 | 12.97 | 2.52 | 0.4 |
| Methanosarcina mazei | | | |
| Replicate 1 | 14.10 | 2.50 | 9.6 |
| Replicate 2 | 13.84 | 2.16 | 9.2 |
| Replicate 3 | 13.71 | 1.95 | 13.1 |
| Replicate 4 | 13.46 | 2.05 | 12.9 |
| Methanobacterium formicicum | | | |
| Replicate 1 | 12.78 | 2.08 | 5.0 |
| Replicate 2 | 8.94 | 1.48 | 2.1 |
| Replicate 3 | 12.75 | 1.80 | 1.6 |
| Replicate 4 | 12.91 | 2.19 | 4.5 |
| Methanobacterium palustre | | | |
| Replicate 1 | 11.93 | 2.22 | 3.7 |
| Replicate 2 | 9.69 | 1.34 | 2.8 |
| Replicate 3 | 10.55 | 1.61 | 2.6 |
| Replicate 4 | 11.30 | 1.53 | 1.4 |
